# Land snail dispersal, abundance and diversity on green roofs

**DOI:** 10.1101/723395

**Authors:** Michael L. McKinney, Nicholas S. Gladstone, Jillian G. Lentz, Faith A. Jackson

## Abstract

We present the first major systematic study of land snail diversity on green roofs. We surveyed 27 green roofs and the adjacent ground habitat in six major cities in the southeastern United States. We found a total of 18 species of land snails, with three considered to be non-native, invasive species. The majority of land snails encountered in surveys are widespread, generalist species, typically adapted to open habitats. Twelve of the land snails encountered are “greenhouse” species that are very commonly transported via the horticultural trade. Therefore, we infer that at least some land snail species are introduced to green roofs via initial green roof installation and associated landscaping. Additionally, some similarity between roof and ground populations indicates dispersal from nearby ground habitats. The major determinants of snail species diversity and community composition are largely derived from local environmental conditions that are significantly correlated to the quality of green roof maintenance regime and plant diversity. Roof area, height, and age are seemingly not significant characteristics that dictate land snail species richness.

## Introduction

Green roofs (i.e., roofs designed to have substrate and vegetation) are increasingly common in many parts of the world. They are an important part of urban green infrastructure, with many environmental benefits relating to storm water runoff, air and water pollution, urban heat island effects, and improved energy conservation [1]. In addition, green roofs can act as vegetated islands of refugia in an otherwise hostile urban matrix to provide habitat for non-human species and promote overall urban biodiversity [2–3]. Previous studies have documented the role of green roofs as habitat for several groups of organisms, especially mobile groups that can readily colonize them, such as birds [4], bees [5], and other major insect groups [6]. These studies document that many native and non-native species can potentially colonize and persist in green roof habitats. However, studies that document green roof colonization and population persistence of animal groups characterized by low vagility (such as land snails) are quite rare. Only one study incorporating land snail surveys on green roofs was located via literature search, documenting four species found on two roofs in Finland [7].

Along with other invertebrate groups ([8] for review), land snails could be common constituents of green roof environments. Generally, most land snail species in North America are associated with moist forest ecosystems, and the potentially harsh conditions of green roof habitats (e.g., prolonged exposure to direct sunlight, comparatively less shelter habitat) may not support diverse land snail communities. Yet, land snails have been well documented in a variety of urban habitats [9–12], are common “hitchhiker” species that are often transported on commercial materials such as horticultural plants [13], and may even disperse via translocation on larger vertebrate animals [14–15]. Moreover, land snails are often seen crawling on building walls and could actively colonize green roofs on their own.

Here, we conduct the first major and systematic survey of snail occupation of green roof habitat. We surveyed 27 green roofs in six major cities in the southeastern United States. Our objectives were to 1) determine how common and how diverse land snails are on green roofs, 2) investigate the relationships between land snail communities and green roof characteristics, and 3) compare land snail diversity of green roofs to immediately adjacent ground habitats.

## Materials and Methods

### Field Sites and Survey Protocol

We identified six major metropolitan areas in the southeastern United States that were known to have buildings with designed green roofs. Three of these cities are in Tennessee: Knoxville, Chattanooga, and Nashville. The other three cities were: Atlanta, Georgia; Charleston, South Carolina, and Savannah, Georgia (Fig 1). Buildings with green roofs were located using several methods: internet searches (e.g., www.greenroofs.com), social media, and contacts with green roof design firms in those cities. Each building with a green roof was then contacted to see if we could gain access to perform a land snail survey. To maximize our sampling diversity, we were interested in green roofs of any size and height.

**Fig 1.**
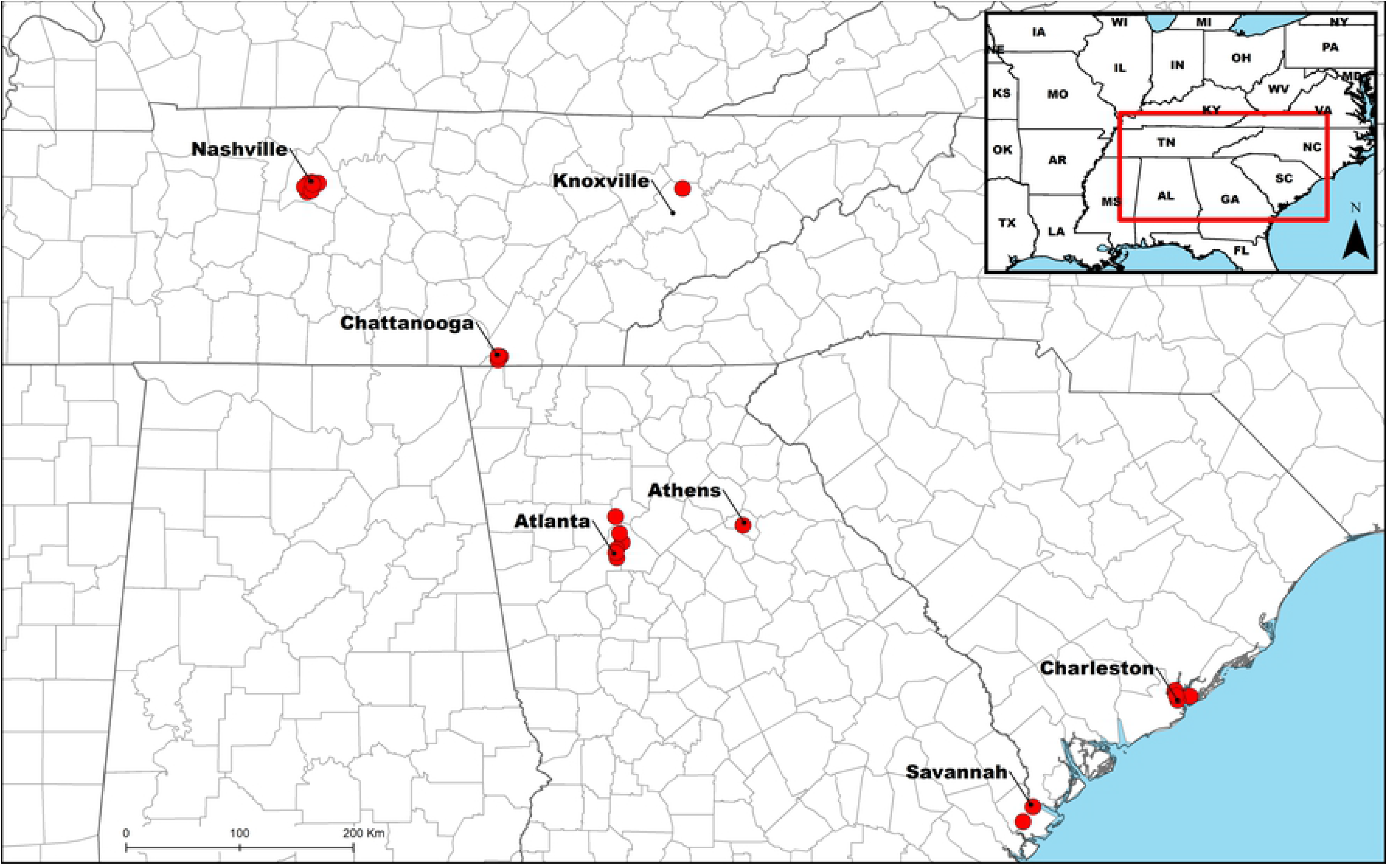
Map of all land snail survey sites with adjacent major cities labeled.

Between March 12, 2018 and November 5, 2018, we visited all identified green roofs that would provide access to our survey team, with a total of 27 sites surveyed. During our visit to each green roof site, we met with facilities managers and collected as much information as possible about each roof: date of construction, green roof installation company, original plant list and details of regular green roof maintenance (irrigation, weeding, plant replacement, etc.). In order to standardize snail sampling effort, we used timed visual searches that were proportional to the area of the roof using the formula of one person searching for 15 minutes per 25 square meters, with no overlap of search areas. During the survey, we collected all dead snail shells that were encountered. Live land snails were photographed in several positions for later identification and then returned to the substrate. Additionally, 0.5 L bags of substrate were collected from each site to search for microsnail species (<5mm in shell diameter; [16]). To standardize substrate sampling effort, one bag of substrate was collected for approximately 25 square meters of roof area. We also collected land snails on the ground around the perimeter of each roof site to examine similarities between roof and ground land snail fauna. We used the same roof sampling procedure for ground collections, with a timed visual search combined with substrate bag samples. Surveys of adjacent ground habitat did not exceed 5 m away from the building.

After the survey, substrate bags were processed with soil sieves (1 micrometer) to extract microsnails and all remaining land snails were identified using published keys and species descriptions [17–19], as well as examination by taxonomic specialists (Dan Dourson). In addition to the land snail surveys, we also noted the dominant vegetation and plant diversity of each green roof. In some cases, this was aided by plant lists provided by the building manager or the company that installed the green roof.

For insight into dispersal modes, we compared our green roof land snail fauna to lists of “greenhouse species” that are commonly transported with plants in the horticultural trade [13]. For insight into adaptive potential to green roof habitats, we searched the internet and literature using various sources for information on the natural habitat where each species is typically found [19–20].

### Data Analyses

Metrics of community diversity and similarity between both roof and ground habitats were quantified for each site, including species richness, total abundance, Simpson’s diversity index (D), Shannon’s index (H’), Evenness (E), and the Jaccard’s index of similarity. To determine whether these metrics were significantly different between roof and ground habitat, we conducted five nested analysis of variance (ANOVA) models using the lme4 package [21]. Habitat type (ground or roof) was utilized as the fixed factor, site location as the random factor, and each diversity metric (richness, abundance, D, H’, E) as a response variable. Residual plots of scaled data were visually assessed and did not reveal any deviations from homoscedasticity or normality. P-values were extracted using the Satterthwaite approximation method with the lmerTest package [22]. To assess variation attributable to fixed and random effects within each model, marginal and conditional R^2^ values were calculated using the MuMIn package [23].

To evaluate the diversity of green roof habitats and their potential relationship to land snail diversity, a dataset containing characteristics relating to each green roof site (roof height, roof area, year of installation, roof type, maintenance and plant diversity) was subjected to principal components analysis (PCA). Data were categorically grouped in relation to snail species richness at each site (low = 0–2 species present, moderate = 3–5, high = 6–9). Data on plant diversity was similarly grouped at each site (low = 0–2 species/m^2^, moderate = 2–4 species / m^2^, high > 4 species / m^2^). For maintenance regime, roofs that had received little or no attention in the preceding year were ranked as low maintenance. Roofs that were watered and weeded at irregular intervals were ranked as moderate maintenance. Roofs that were watered and weeded on a regular basis at least every 3 months were ranked as high maintenance. Correlation coefficients (PC loadings) of each variable were assessed to ascertain the significance of green roof traits to each PC. PC scores of the 27 green roofs were correlated with species richness using a Spearman Rank Correlation (SRC) analysis. Additionally, to investigate if faunal similarity of the ground and roof habitats were related to green roof traits, the Jaccard’s index of similarity was also subject to SRC with each green roof trait.

Lastly, to further investigate the relationship between green roof characteristics and land snail diversity on green roofs, a separate species diversity matrix was generated and subsequently scaled to minimize influence of considerably abundant groups. Scaled data was converted to a Bray-Curtis dissimilarity matrix and subject to permutational multivariate analysis of variance (PERMANOVA) to test the effect of green roof traits on species richness. This test was performed in the vegan package in R for 999 permutations [24]. P-values obtained from pairwise comparisons were corrected using a Bonferroni test.

## Results

### Survey Results and Species Composition

A total of 27 green roof sites were analyzed, ranging from 5–191,000 sq. ft, from 2–53 years old, and up to 8 stories high (Table 1). Plant diversity on these roofs was roughly evenly distributed between low, medium and high, with *Sedum* being the dominant vegetation on many roofs (S1 Table) as is commonly the case with green roofs due to their drought tolerance. Land snail species richness ranged from zero to nine species, with seven sites having no species observed and only five surveyed sites containing more than three species.

**Table 1.**
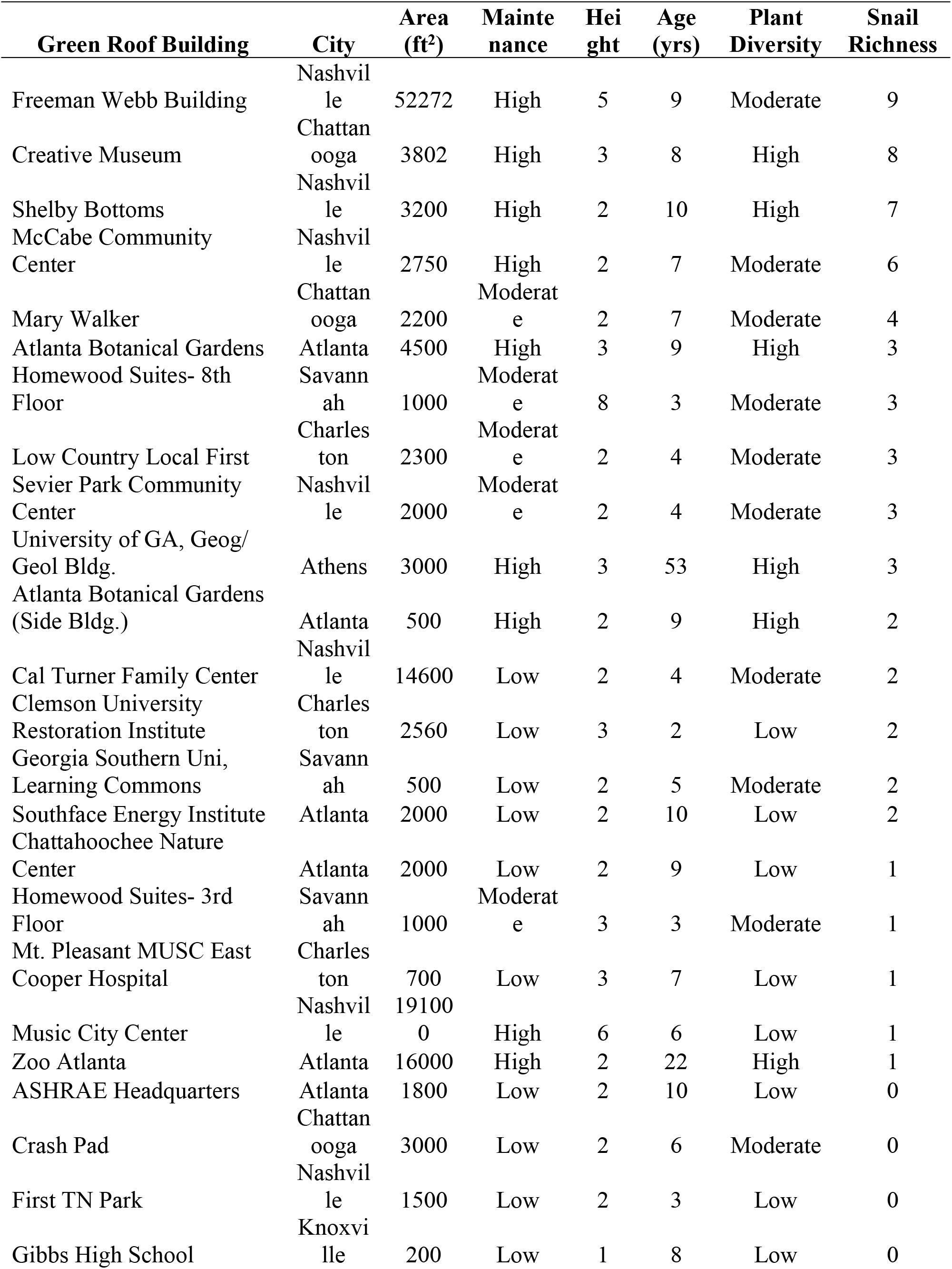
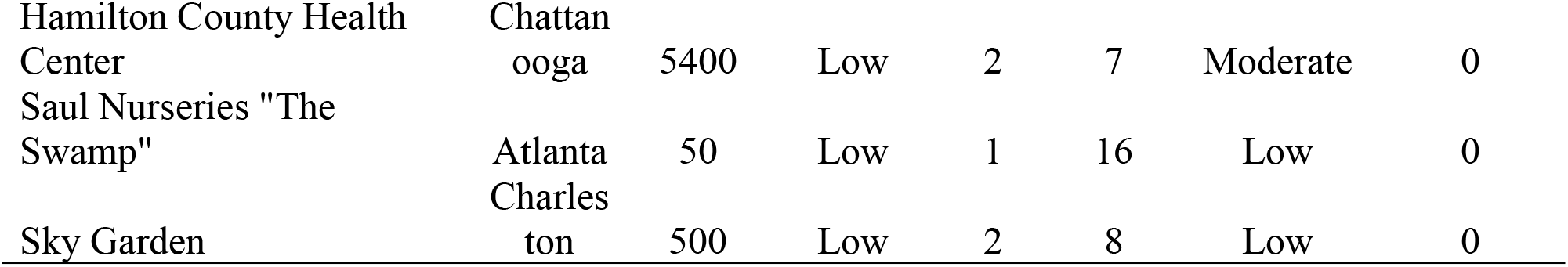
Summary of green roof survey results ordered by snail richness.

A total of 18 land snail species were found on the 27 green roofs with the taxa found on the most green roof sites being: *Succinea sp., Zonitoides arboreus, Polygyra cereolus*,and *Pupoides albilabis* (Fig 2), each being found on seven or more sites. These four species were also the most abundant with a total of 61, 126, 90 and 64 individuals of each being found, respectively (Table 2). Thus, these four species account for 73.2% of all land snails encountered on green roofs. A substantial number of snails found showed evidence of a living population: 44 of 67 occurrences (67.7%) contained at least some snails that were recently dead (with tissue) or still alive (S1 Table).

**Fig 2.**
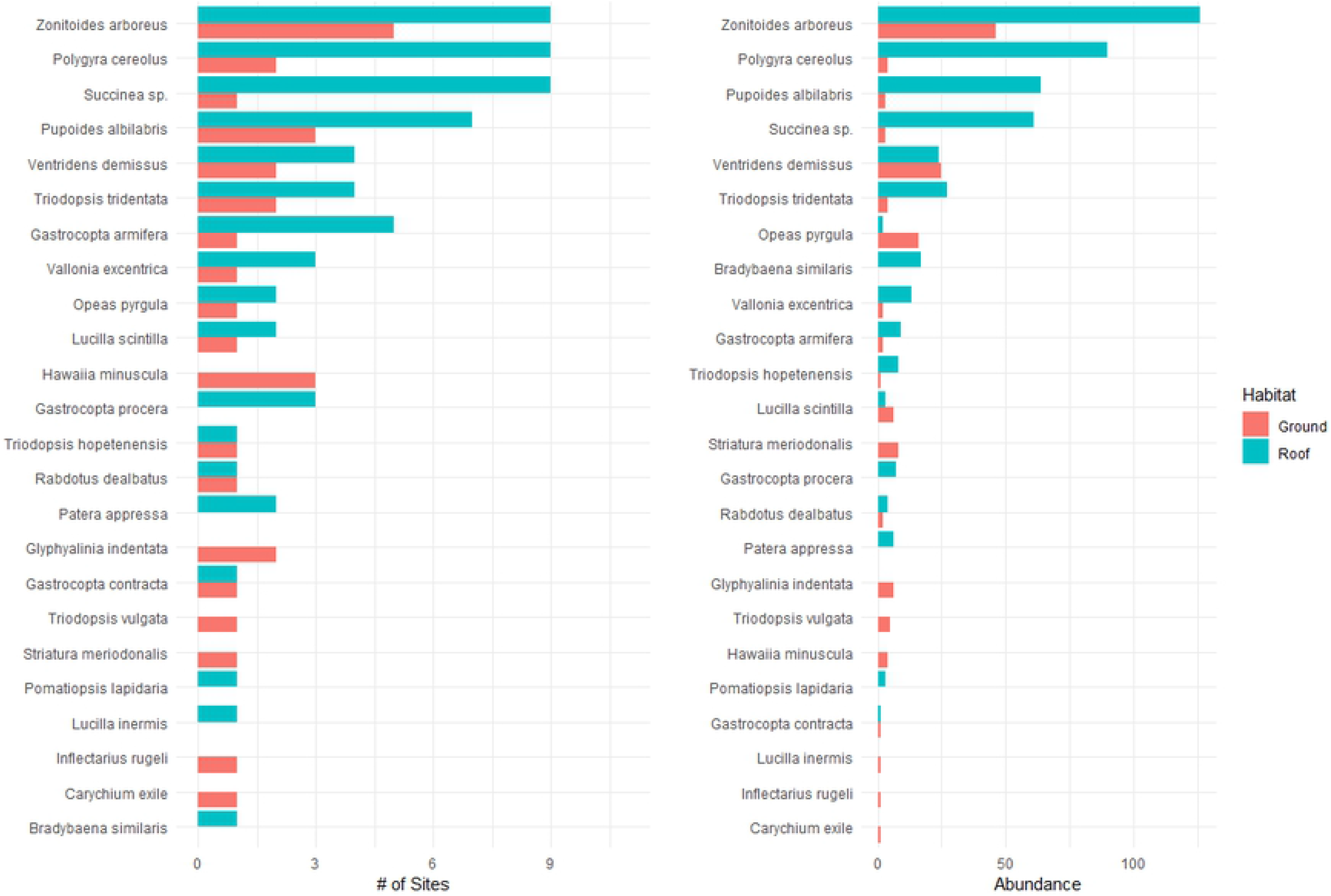
Snail diversity summarizing number of green roof sites found and total abundance.

**Table 2.**
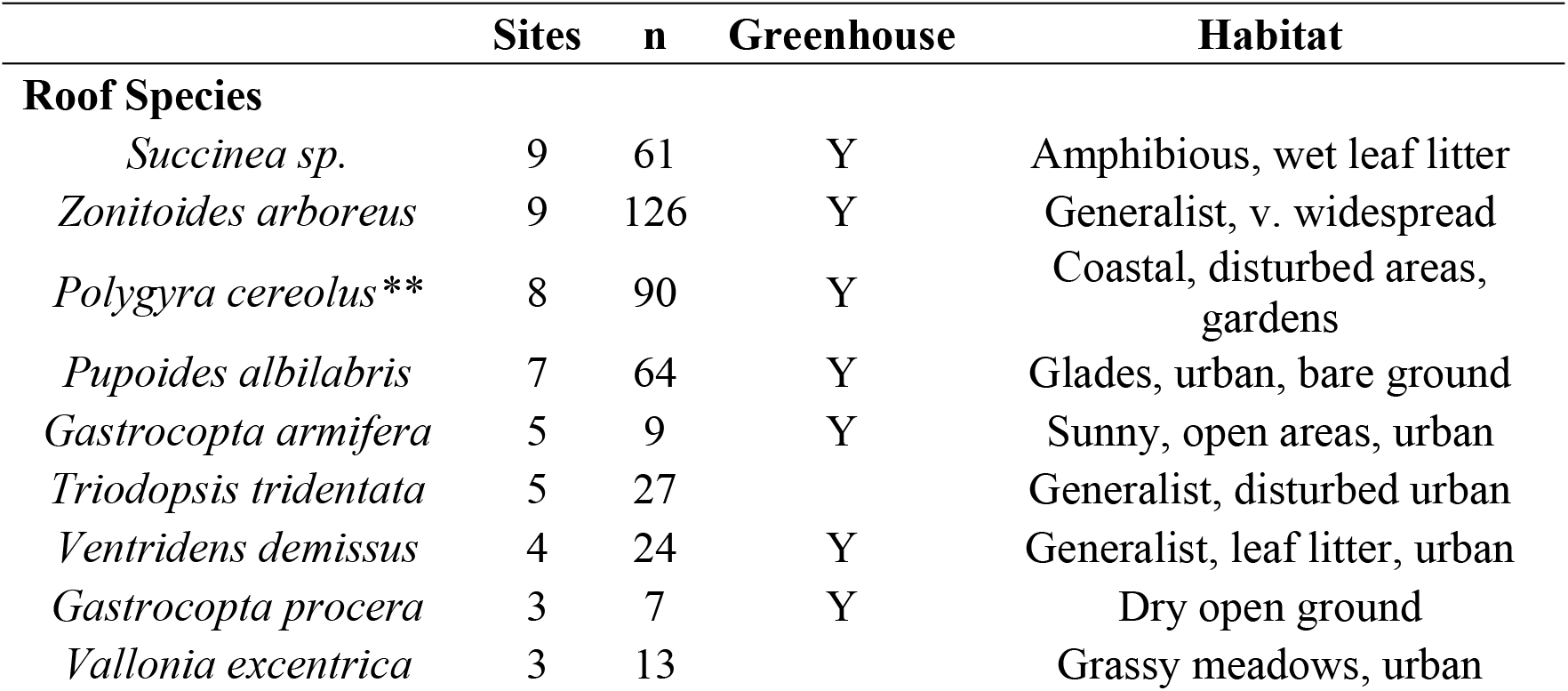
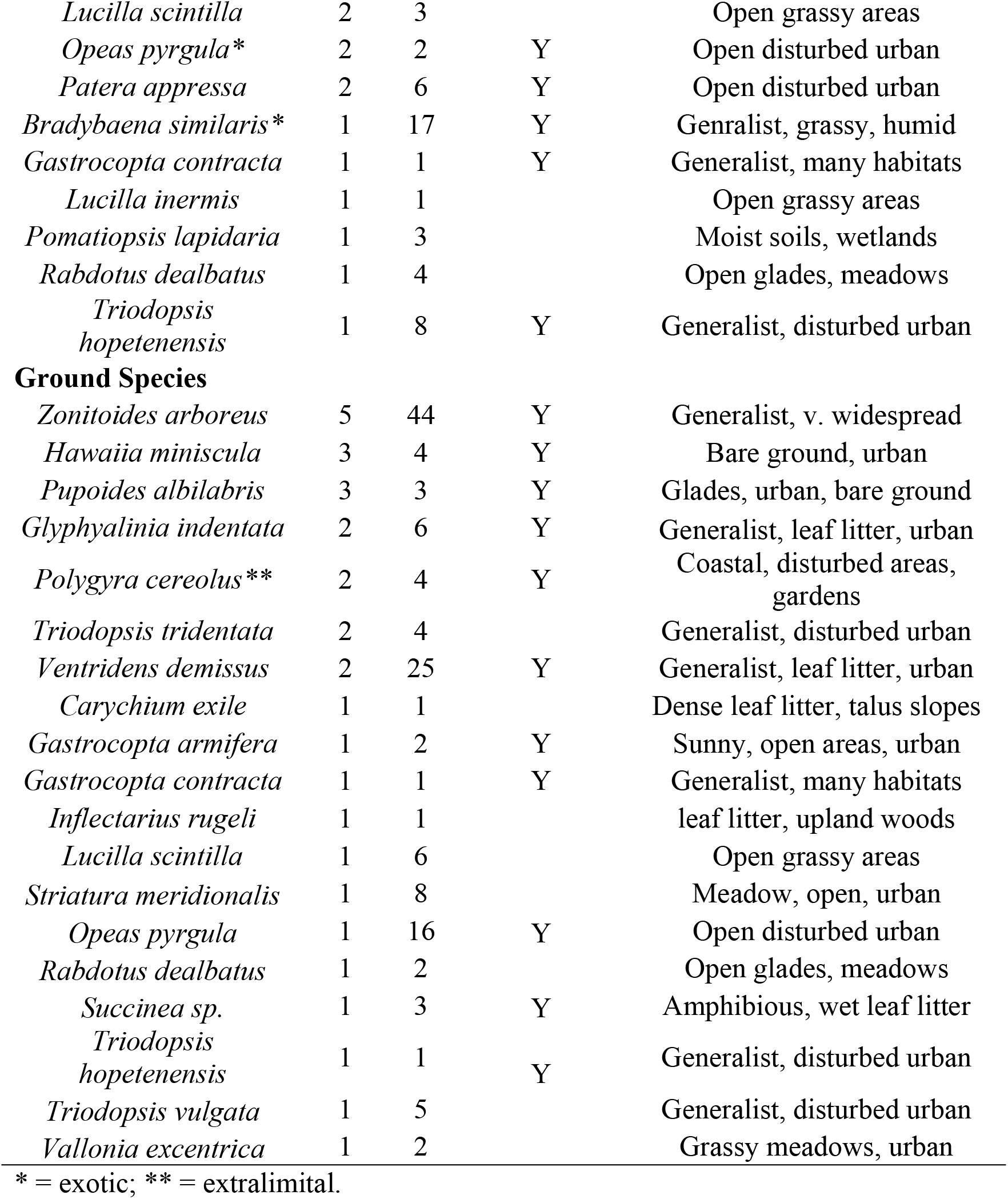
Summary of green roof and ground-dwelling species. “Greenhouse” designation in relation to survey results from [13]. Habitat information was obtained from [19–20].

Interestingly, twelve of the 18 green roof land snail species are common “hitchhiker” species (Table 2) found in greenhouses which serve as the transport hubs of horticultural plant species [13]. The general ecological adaptations and habitat preferences of the 18 species of green roof land snails are similar, with most, if not all of them, being generalists adapted to open, grassy and/or disturbed anthropogenic habitats [17, 19]. However, as discussed below there are differences in the maintenance regimes of these 27 roofs that affect their habitability and snail fauna.

Only three of the 18 green roof species are not native to their locations. We found two exotic species (*Opeas pyrgula*, *Bradybaena similaris*) native to southeastern Asia and one known extralimital species (*Polygyra cereolus*). The most widespread and abundant of these three non-natives was *Polygyra cereolus*, with 90 individuals found at eight green roofs in five cities. *Opeas pyrgula* was found at two green roofs in two cities (one individual each) and *Bradybaena similaris* had 17 individuals at only one location. In addition to these three species, it is possible that at least some of the succinids are not native but this is uncertain due to the difficulty in differentiating species in this taxonomically challenging group [13].

On the ground habitat adjacent to the green roofs, we found a total of 19 species (Table 2). Two of the most abundant species (*Zonitoides arboreus*, *Pupoides albilabis*) were also the among the most abundant on green roofs. Six (31.6%) of these ground species were not found in the pool of green roof species. Ten (52.6%) of ground species are listed as hitchhiker, greenhouse species compared to twelve (66.7%) of roof species. We found the same species of non-natives on the ground as on the green roofs (*Opeas pyrgula, Polygyra cereolus*) apart from *Bradybaena similaris* which was absent from all ground samples.

### Data Analyses

Nested ANOVA results show a significant difference between green roof and ground habitats in species richness, Simpson’s index, and Evenness (Table 3). Site location accounted for the vast majority of variation in the dataset, indicating that individual roof and ground diversity are likely location dependent. In most cases, we were not able to generate a value for the Jaccard Index, as there were many sites with no land snails on the roof, on the ground, or both. However, the three sites with the highest roof species richness (Freeman Webb Building, Creative Museum, Shelby Bottoms) had a Jaccard Index of 0.385, 0.125, and 0.125 respectively. The highest overall Jaccard Index of all sites was 0.5 at the McCabe Community Center with six species on both the roof and ground habitat of which three species were shared. These results indicate that, even in instances of having viable roof and ground habitat for land snails, species composition is often not similar between the two habitats.

**Table 3.** Nested ANOVA results.

| Response | F | $R^2m$ | $R^2c$ | P |
| --- | --- | --- | --- | --- |
| Richness | 4.110 | 0.054 | 0.656 | 0.026* |
| Abundance | 2.743 | 0.086 | 0.189 | 0.079 |
| D | 3.736 | 0.097 | 0.358 | 0.035* |
| H' | 2.805 | 0.045 | 0.588 | 0.076 |
| E | 3.7262 | 0.085 | 0.486 | 0.035* |

\*significant p-value

The first three principal components (PCs) account for 80.15% of the total variation for the green roof trait dataset (Fig 3). PC 1 (37.57%) was primarily influenced by the positive loadings of roof maintenance (0.5571) and plant diversity (0.5803). PC 2 (28.27%) was primarily influenced by the negative loadings of roof height (−0.6471) and roof area (−0.6661), and PC 3 (14.34%) was primarily influenced by type of green roof (0.390). SRC test showed a significant correlation between species richness with resulting PC scores, and faunal similarity (Jaccard’s index) between roof and ground habitat was not significantly related to any individual green roof characteristic. Moreover, additional SRC tests show that roof species richness is significantly correlated with both roof maintenance and plant diversity. However, subsequent PERMANOVA results show no significant relationship between species richness and any green roof characteristic. This may, in part, be due to the substantial amount of previously revealed location-dependency.

**Fig 3.**
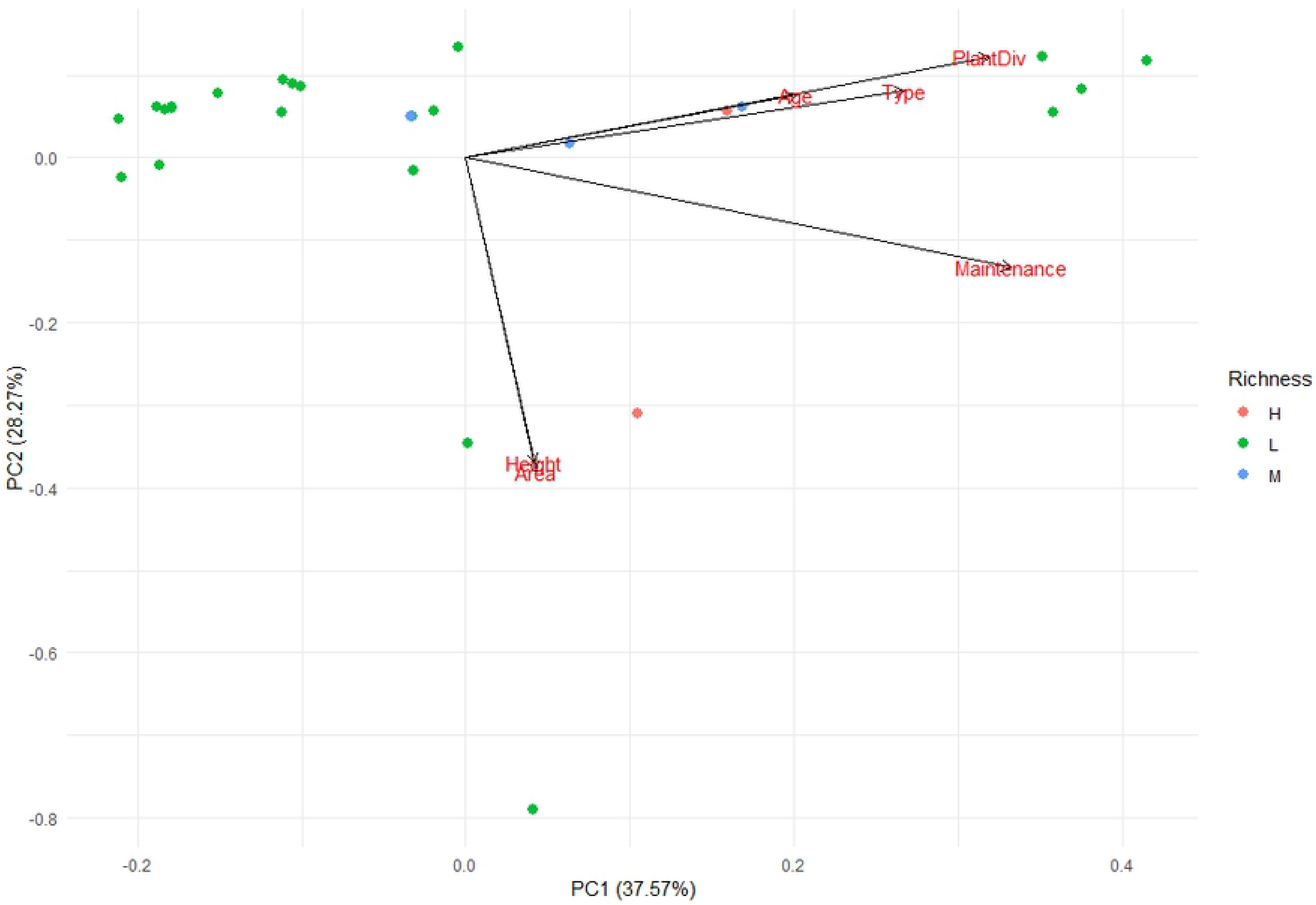
PCA results showing each of the six green roof characteristic loadings. Green roofs are represented by individual points and categorized by level of species richness (low, medium, high).

## Discussion

### Snail Diversity and Environmental Influences

We found a considerable abundance (466) and species richness (18) of land snails on green roofs. However, it is notable that over one-fourth (25.9%) of the green roofs had no land snails. These snail-free green roofs were all categorized as low-maintenance, including some green roofs that had experienced no maintenance (watering, weeding, replanting) for years (e.g., Gibbs High School). In addition, most of these snail-free, low maintenance green roofs also had low plant diversity. Conversely, the roofs that were most species-rich (with three or more land snail species) all had moderate-high maintenance regimes and also had moderate-high plant diversity. These observations imply that increasing land snail species richness is at least somewhat related to increasing green roof maintenance and plant diversity. This makes sense given that some of the major habitat variables that promote land snail diversity and abundance are moisture, vegetative cover and coarse woody debris [12, 25–26]. A completely neglected green roof with little or no watering during drought periods and sparse vegetation will likely be poor habitat for land snails, especially in the hot summer months of the southeastern U.S.

On the other hand, although there were significant correlations between species richness and green roof characteristics, the multivariate analyses indicates that species richness is not readily explained by any single variable. In general, we find that land snail diversity and abundance is best explained by somewhat idiosyncratic local environmental conditions. Interestingly, this dominant effect of local environmental conditions on diversity had also been found in green roof studies of other taxa such as beetles, where local habitat traits have a stronger effect on community composition than landscape variables [27]. This local effect seems to be especially strong for low-mobility invertebrates [28]. An example of this location-dependency in our data is the green roof at Zoo Atlanta – one of the oldest (22 years), largest, highly vegetated, and well-maintained sites. Despite these characteristics, only one snail species was discovered – a very common ground-dwelling species at the zoo (*Ventridens demissus*). Because of this anomalous finding, a second roof survey was done at this location and confirmed this low diversity. Follow-up conservations with the Zoo Atlanta green roof management indicated that this anomalously low snail diversity may be attributable to the fact that no new horticultural plants have been added to the roof since the green roof installation 22 years ago. If horticultural transport is a major source of dispersal, as noted below, then it may be that snail populations on green roofs may become extirpated if not regularly replaced.

### Dispersal to Roofs

Most studies of green roof faunal colonization look at highly mobile taxa such as flying insects which can readily colonize green roof habitat via their own locomotion [2, 8]. For organisms characterized by low vagility, an alternative mechanism of dispersal (i.e., translocation via landscaping and horticulture) may be the most influential for colonizing green roofs. It is well documented that plant nurseries are hot spots for many land snail species and that horticultural plants are major mechanisms of land snail introductions [13, 29]. In this study, we find that most land snail species encountered are indeed well-documented greenhouse inhabitants (Table 2). A similar mechanism has been suggested for the dispersal of Collembola (springtails) onto green roofs via composting for roof soil enrichment [30]. If our findings are validated with further research, it could provide useful insight into the dynamics of how green roof ecosystems are created and change through time. It also indicates that humans can have some control over which land snails colonize these designed ecosystems.

In addition to hitchhiking, our findings may also imply a role for self-dispersal via land snail locomotion. Snail locomotion is famously slow. For example, Bergey [31] recently documented that the spread of the non-native *Cornu aspersum* across 16 residential yards in a single city block in Norman, Oklahoma took six years. The presence of vegetation adjacent to many buildings, and the common tendency of land snails to crawl on the (often calcium-rich) walls of such buildings, indicates that land snails could likely colonize green roofs under the right conditions and given enough time. The two most species-rich roofs (Freeman Webb building and the McCabe Community Center) and their nearby ground habitats share several species that could indeed indicate active land snail dispersal and interchange between ground and roof communities. Both green roofs are well maintained and have a moderate plant diversity composed of quite a few native plants, ostensibly increasing their habitability for land snails. However, the majority of our findings show that there is generally little similarity between the pool of roof and ground species. Given that environmental conditions of green roofs are often quite different (e.g., much hotter and drier) from nearby ground conditions, different abiotic tolerances may be required to persist in each respective habitat. This is especially true for green roofs that are not well maintained and/or have very low plant diversity and vegetative cover (e.g., *Sedum* monocultures). Moreover, it is important to note that many ground habitats adjacent to our green roof sites are mainly “hardscapes” with little or no vegetation. This is reflected in our survey data which often found no ground snails (S2 Table) and thus reduces our sample size to make roof-ground comparisons.

### Comparison to Other Green Roof Fauna

As has been found with several other invertebrate taxa [2, 27, 32], the species inhabiting green roofs tend to be widespread, generalist and disturbance-adapted species. The same studies also show that just a few species tend to dominate in terms of total abundance and being the most widespread among green roof habitats. This is true in our study where just four species account for almost three-fourths of all snails found. Also as found in many other invertebrate studies [5, 32), most of our green roof species are native, with a minority (at least three species) being invasive, broadly adapted non-native species.

In terms of species richness, our findings indicate that land snails on green roofs may be considerably less diverse than more mobile taxa. For example, Kratschmer and others [5] surveyed just nine green roofs in Vienna, Austria, and identified 90 wild bee species. A study of 17 green roofs in Switzerland found 161 highly dispersive beetle species [27]. On the other hand, our findings are similar for non-flying taxa. A review of 102 green roofs in Switzerland found just 91 ground beetle species [32].

## Conclusions and Practical Applications

In their review of invertebrates on green roofs, MacIvor and Ksiazek [8] point out that “it is not clear whether they adequately provide habitat or not”. They also note that green roof habitats can vary widely, with some green roofs providing very little habitat for invertebrates. Their relative isolation in a hostile urban matrix prevents colonization and this often combines with the harsh conditions on roof tops to inhibit long-term population persistence. Therefore, we need many studies of all major invertebrate groups to truly understand the extent that green roofs can promote invertebrate biodiversity in urban areas. While several invertebrate taxa have seen a growing literature on green roof habitats, here we provide the first extensive study of green roof habitat for land snails.

Our study indicates that well maintained green roofs can support a modest land snail community of mostly native taxa. Moreover, our findings further support the importance of local environmental controls on diversity and community composition in green roof ecosystems. Habitat quality clearly varies with key factors such as plant diversity and maintenance regimes. This explains why attempts to find simple island biogeography relationships to green roof data have not been productive [33]. Island biogeographic factors such as area, building height (distance to ground), and distance between roofs are generally less important determinants of diversity and community composition than habitat quality (which can vary widely). Furthermore, if dispersal is often driven by direct human introduction by transport of cultivated plants then dispersal factors such as distance to source populations in nearby ground habitats become irrelevant.

## Acknowledgements

We thank all of the property and business owners that allowed us to survey the green roofs, the maintenance workers that gave us additional information for each site, and the landscaping workers that gave us direction for potential site selection. Additionally, we thank Dan Dourson for help in identifying land snail specimens.

## References

1. Cascone S. Green roof design: state of the art on technology and materials. Sustainability 2019;11(11): 3020.

2. Williams NSG, Lundholm J, MacIvor SJ. Do green roofs help urban biodiversity conservation?. J Appl Ecol 2014;51(6): 1643–1649.

3. Yildirim S, Ozden O. Positive effects of vegetation: Biodiversity and extensive green roofs for Mediterranean climate. International Journal of Advanced and Applied Sciences 2018;5(10) 87–92.

4. Fernández CR, Redondo PG. Green roofs as a habitat for birds: a review. J Anim Vet Adv 2010;9(15): 2041–2052.

5. Kratschmer S, Kriechbaum M, Pachinger B. Buzzing on top: Linking wild bee diversity, abundance and traits with green roof qualities. Urban Ecosyst 2018;21(3): 429–446.

6. MacIvor SJ, Lundholm J. Insect species composition and diversity on intensive green roofs and adjacent level-ground habitats. Urban Ecosyst 2011;14(2): 225–241.

7. Páll-Gergely B, Kyrö K, Lehvävirta S, Vilisics F. Green roofs provide habitat for the rare snail (Mollusca, Gastropoda) species *Pseudotrichia rubiginosa* and *Succinella oblonga* in Finland. Memoranda Societatis pro Fauna et Flora Fennica 2014;90: 13–15.

8. MacIvor SJ, Ksiazek K. Invertebrates on green roofs. In: Sutton RK, editor. Green roof ecosystems. Springer, Cham, 2015. pp. 333–355.

9. Lososová Z, Horsák M, Chytrý M, Čejka T, Danihelka J, Fajmon K, et al. Diversity of Central European urban biota: effects of human‐made habitat types on plants and land snails. J Biogeogr 2011;38(6): 1152–1163.

10. Horsák M, Čejka T, Juřičková L, Wiese V, Horsáková V, Lososová Z. Drivers of Central European urban land snail faunas: the role of climate and local species pool in the representation of native and non-native species. Biol Invasions 2016;18(12): 3547–3560.

11. Barbato D, Benocci A, Caruso T, Manganelli G. The role of dispersal and local environment in urban land snail assemblages: an example of three cities in Central Italy. Urban Ecosyst 2017;20(4): 919–931.

12. Hodges MN, McKinney ML. Urbanization impacts on land snail community composition. Urban Ecosyst 2018;21(4): 721–735.

13. Bergey EA., Figueroa LL, Mather CM, Martin RJ, Ray EJ, Kurien JT, et al. Trading in snails: plant nurseries as transport hubs for non-native species. Biol Invasions 2014;16(7): 1441–1451.

14. Rees WJ. The aerial dispersal of Mollusca. J Molluscan Stud 1965;36(5): 269–282.

15. Wada S, Kawakami K, Chiba S. Snails can survive passage through a bird’s digestive system. J Biogeogr 2012;39(1): 69–73.

16. Liew T, Clements R, Schilthuizen M. Sampling micromolluscs in tropical forests: one size does not fit all. Zoosymposia 2008;1: 271–280.

17. Pilsbry HA. Land Mollusca of North America (north of Mexico). Monograph 3. Academy of Natural Sciences of Philadelphia 2(2); 1948.

18. Burch JB. How to know the eastern land snails. No. 594.3, B87; 1962.

19. Dourson DC, Burch JB, Emberton KC, Forsyth R, Karstad A, Osborne BJ, et al. Kentucky’s land snails and their ecological communities. Goatslug Publications; 2010.

20. Hubricht L. The distributions of the native land mollusks of the eastern United States. Chicago: Field Museum of Natural History; 1985.

21. Bates D, Sarkar D, Bates MD, Matrix L. The lme4 package. R package version 2007;2(1): 74.

22. Kuznetsova A, Brockhoff PB, Christensen RGB. lmerTest package: tests in linear mixed effects models. Journal of Statistical Software 2017;82(13).

23. Barton K. MuMIn: multi-model inference. R package version 1. 0. 0. 2009; http://r-forge.r-project.org/projects/mumin/

24. Oksanen J, Blanchet FG, Kindt R, Legendre P, O’hara RB, Simpson GL, et al. Vegan: community ecology package. R package version 1.17-4. 2010; http://cran.r-project.org>. Acesso em 23

25. Kappes H, Kopeć D, Sulikowska-Drozd A. Influence of habitat structure and conditions in floodplain forests on mollusc assemblages. Pol J Ecol 2014;62(4): 739–751.

26. Kirchenbaur T, Fartmann T, Bässler C, Löffler F, Müller J, Strätz C, et al. Small-scale positive response of terrestrial gastropods to dead-wood addition is mediated by canopy openness. For Ecol Manage 2017;396: 85–90.

27. Kyrö K, Brenneisen S, Kotze JD, Szallies A, Gerner M, Lehvävirta S. Local habitat characteristics have a stronger effect than the surrounding urban landscape on beetle communities on green roofs. Urban For Urban Green 2018; 29: 122–130.

28. Braaker S, Ghazoul J, Obrist MK, Moretti M. Habitat connectivity shapes urban arthropod communities: the key role of green roofs. Ecology 2014;95(4): 1010–1021.

29. Cowie RH, Hayes KA, Tran CT, Meyer III WM. The horticultural industry as a vector of alien snails and slugs: widespread invasions in Hawaii. Int J Pest Manag 2008;54(4): 267–276.

30. Joimel S, Grard B, Auclerc A, Hedde M, Le Doaré N, Salmon S, et al. Are Collembola “flying” onto green roofs?. Ecol Eng 2018;111: 117–124.

31. Bergey EA. Dispersal of a non-native land snail across a residential area is modified by yard management and movement barriers. Urban Ecosys 2019;22(2): 325–334.

32. Pétremand G, Chittaro Y, Braaker S, Brenneisen S, Gerner M, Obrist MK, et al. Ground beetle (Coleoptera: Carabidae) communities on green roofs in Switzerland: synthesis and perspectives. Urban Ecosyst 2018;21(1): 119–132.

33. Blank L, Vasl A, Schindler BY, Kadas GJ, Blaustein L. Horizontal and vertical island biogeography of arthropods on green roofs: a review. Urban Ecosyst 2017;20(4): 911–917.

